# Comparative analysis of transcriptomic and proteomic expression between two non-small cell lung cancer subtypes

**DOI:** 10.1101/2024.09.05.611373

**Authors:** Ben Nicholas, Alistair Bailey, Katy J McCann, Peter Johnson, Tim Elliott, Christian Ottensmeier, Paul Skipp

## Abstract

Non-small cell lung cancer (NSCLC) is frequently diagnosed at an advanced stage and has poor survival. NSCLC subtypes require different treatment regimes, hence there are extensive efforts to find more precise and non-invasive differential diagnostics tools. Complementing these efforts, we examined two NSCLC subtypes for differences that may inform treatment options and identify potential novel therapeutic pathways.

Here we present a comparative analysis of transcriptomic and proteomic expression in tumours from a cohort of 22 NSCLC patients: 8 squamous cell carcinoma (LUSC), 14 adenocarcinoma (LUAD). We examined differential gene and differential protein expression between LUSC and LUAD, and between NSCLC subtypes and either PBMCs or normal adjacent lung tissue (NAT).

We found that both NSCLC subtypes shared common differences in gene expression to PBMC relating to developmental and structural changes, and common protein expression differences to NAT relating to protein translation and RNA related processing and splicing.

Between NSCLC subtypes we found differential gene expression relating to cell differentiation for LUSC and cellular structure and immune response regulation for LUAD. Differential protein expression between NSCLC subtypes related to extracellular structure for LUSC and metabolic processes, including glucose metabolism for LUAD.

Many of our observations of differentially expressed genes and proteins between NSCLC subtypes support and inform existing observations, aiding both basic and clinical research seeking to identify subtype biomarkers or druggable targets.

## Introduction

Lung cancer is the second most common cancer in the UK, with a majority of cases diagnosed at advanced stages, either locally advanced (stage III) or metastatic (stage IV). Non-small cell lung cancer (NSCLC) comprises 85-90% of these cases and is further categorised into three histological subtypes: adenocarcinoma (LUAD), the most common type, typically develops in the alveoli of the outer peripheral lung. Squamous cell carcinoma (LUSC), the second most frequent type, usually forms in squamous cells located more centrally in the lungs. Large cell undifferentiated carcinoma, the least common, can originate anywhere in the lung [1]. In the UK, less than 20% of all lung cancer patients survive for 5 years, with the majority of patients surviving less than one year post-diagnosis [2,3].

Consequently, there are extensive efforts to find more precise and non-invasive diagnostics for NSCLC, for example using circulating proteins [4] or miRNAs [5]. Likewise, the longitudinal NSCLC TRACERx (TRAcking Cancer Evolution through therapy (Rx)) study has sought to identify the evolutionary processes that help explain disease progression and treatment resistance [6]. Furthermore, we have previously presented identification of HLA presented neoantigens as cancer vaccine targets in two NSCLC subtypes, squamous cell carcinoma (LUSC) and adenocarcinoma (LUAD) [7].

Here we present a comparative analysis of transcriptomic and proteomic expression in tumours from a cohort of 22 NSCLC patients: 8 LUSC and 14 LUAD. The patients were from the same cohort as for our neoantigen study. Using RNA sequencing (RNAseq) and label free quantification (LFQ) of bottom-up mass spectrometery proteomics, we sought to identify differences that may inform treatment options and therapeutic pathways. NSCLC subtype transcriptomes were compared to each other, and to peripheral blood mononuclear cells (PBMC). NSCLC subtype proteomes were also compared to each other and to normal adjacent lung tissue (NAT).

Using differential expression analysis, we identified genes and proteins that characterise each NSCLC subtype.

Our observations offer independent corroboration and contrast to existing studies to aid further research to identify NSCLC subtype biomarkers or targets for more effective subtype specific treatments.

## Results

### NSCLC patient cohort

Table 1 summaries our cohort of 22 NSCLC patients with either LUSC (n=8) or LUAD (n=14) subtype. Tumour tissues underwent RNAseq and mass spectrometry proteomics LFQ. Whole exome sequencing (WES) was used to calculate tumour purity and ploidy [7–9].

**Table 1:** Summary of patients in this study with non-small cell lung cancer

| Donor | Cancer subtype | Smoking status | Tumour wet weight (mg) | Tumour purity | Tumour ploidy | *TIL status |
| --- | --- | --- | --- | --- | --- | --- |
| A113 | LUSC | Current smoker | 26 | 1.0 | 2.0 | Mod |
| A115 | LUSC | Ex smoker | 129 | 0.5 | 1.9 | Mod |
| A116 | LUSC | Never smoker | 113 | 0.5 | 2.5 | Low |
| A133 | LUSC | Ex smoker | 85 | 1.0 | 4.3 | High |
| A134 | LUSC | Current smoker | 152 | 0.5 | 1.8 | High |
| A140 | LUSC | Ex smoker | 71 | 0.4 | 2.7 | High |
| A144 | LUSC | Ex smoker | 122 | 0.5 | 3.8 | High |
| A152 | LUSC | Ex smoker | 288 | 0.3 | 4.7 | - |
| A114 | LUAD | Ex smoker | 1,085 | 1.0 | 2.0 | Mod |
| A117 | LUAD | Current smoker | 158 | 0.3 | 1.9 | Low |
| A118 | LUAD | Ex smoker | 132 | 1.0 | 2.0 | Low |
| A120 | LUAD | Ex smoker | 18 | 1.0 | 2.0 | Mod |
| A136 | LUAD | Never smoker | 98 | 0.3 | 2.0 | High |
| A137 | LUAD | Ex smoker | 270 | 1.0 | 2.0 | High |
| A139 | LUAD | Ex smoker | 93 | 0.4 | 1.6 | Mod |
| A141 | LUAD | Ex smoker | 391 | 0.3 | 5.2 | Mod |
| A142 | LUAD | Current smoker | 277 | 1.0 | 2.6 | High |
| A143 | LUAD | Ex smoker | 97 | 0.5 | 1.9 | Mod |
| A146 | LUAD | Current smoker | 99 | 1.0 | 2.0 | High |
| A147 | LUAD | Current smoker | 88 | 1.0 | 2.2 | High |
| A148 | LUAD | Ex smoker | 229 | 1.0 | 2.0 | High |
| A153 | LUAD | Ex smoker | 436 | 0.4 | 2.6 | - |

\*Tumour-infiltrating lymphocytes

PBMCs were available for RNAseq for 10 of the LUAD patients and 5 of of the LUSC patients. 9 LUAD and 5 LUSC patients had NAT available for proteomics analysis (Table 2).

**Table 2:** Summary of sample availability for patients in this study with non-small cell lung cancer

| Donor | Cancer subtype | Tumour RNAseq | ^PBMC RNAseq | Tumour proteome | *NAT proteome |
| --- | --- | --- | --- | --- | --- |
| A113 | LUSC | Yes | No | Yes | No |
| A115 | LUSC | Yes | No | Yes | No |
| A116 | LUSC | Yes | No | Yes | No |
| A133 | LUSC | Yes | Yes | Yes | Yes |
| A134 | LUSC | Yes | Yes | Yes | Yes |
| A140 | LUSC | Yes | Yes | Yes | Yes |
| A144 | LUSC | Yes | Yes | Yes | Yes |
| A152 | LUSC | Yes | Yes | Yes | Yes |
| A114 | LUAD | Yes | No | Yes | No |
| A117 | LUAD | Yes | No | Yes | No |
| A118 | LUAD | Yes | No | Yes | No |
| A120 | LUAD | Yes | No | Yes | No |
| A136 | LUAD | Yes | Yes | Yes | Yes |
| A137 | LUAD | Yes | Yes | Yes | Yes |
| A139 | LUAD | Yes | Yes | Yes | Yes |
| A141 | LUAD | Yes | Yes | Yes | Yes |
| A142 | LUAD | Yes | Yes | Yes | No |
| A143 | LUAD | Yes | Yes | Yes | Yes |
| A146 | LUAD | Yes | Yes | Yes | Yes |
| A147 | LUAD | Yes | Yes | Yes | Yes |
| A148 | LUAD | Yes | Yes | Yes | Yes |
| A153 | LUAD | Yes | Yes | Yes | Yes |

^Peripheral blood mononuclear cells, \*Normal adjacent tissue

Although it is not technically correct to describe genes or proteins as expressed, transcripts are expressed and proteins are the products of translation, the word expression has become synonymous for the product of a biological process. Hence, we here refer throughout to the quantification of transcripts and peptides as gene expression and protein expression respectively [10,11].

For all 22 NSCLC patients we calculated differential gene expression (DEG) and differential protein expression (DEP) between LUSC and LUAD. We calculated DEG between LUSC and PBMC (n=5) and DEP between LUSC and NAT (n=5). Likewise we calculated DEG between LUAD and PBMC (n=10) and DEP between LUAD and NAT (n=9).

### Comparison of the transcriptomes

As detailed in the materials and methods, count matrices for each sample were calculated containing gene expression values as represented by transcript abundance counts for each gene. One matrix was calculated from genomic alignments and feature counting [12,13] and a second matrix from transcript classification [14] (Tables S1-6).

To examine how well NSCLC subtypes cluster, within-group similarity and to identify batch effects or outlier individuals we performed Principal Component Analysis (PCA) of the normalised feature count data for the top 500 genes with highest variability across samples [15] (Figure 1). The comparison between tumour and PBMC shows clear separation along PC1 with very close clustering of the PBMC samples (Figure 1 A-B). Both the LUSC and LUAD samples were slightly more spread out along PC2, but otherwise grouped together (Figure 1 A-B).

**Figure 1:**
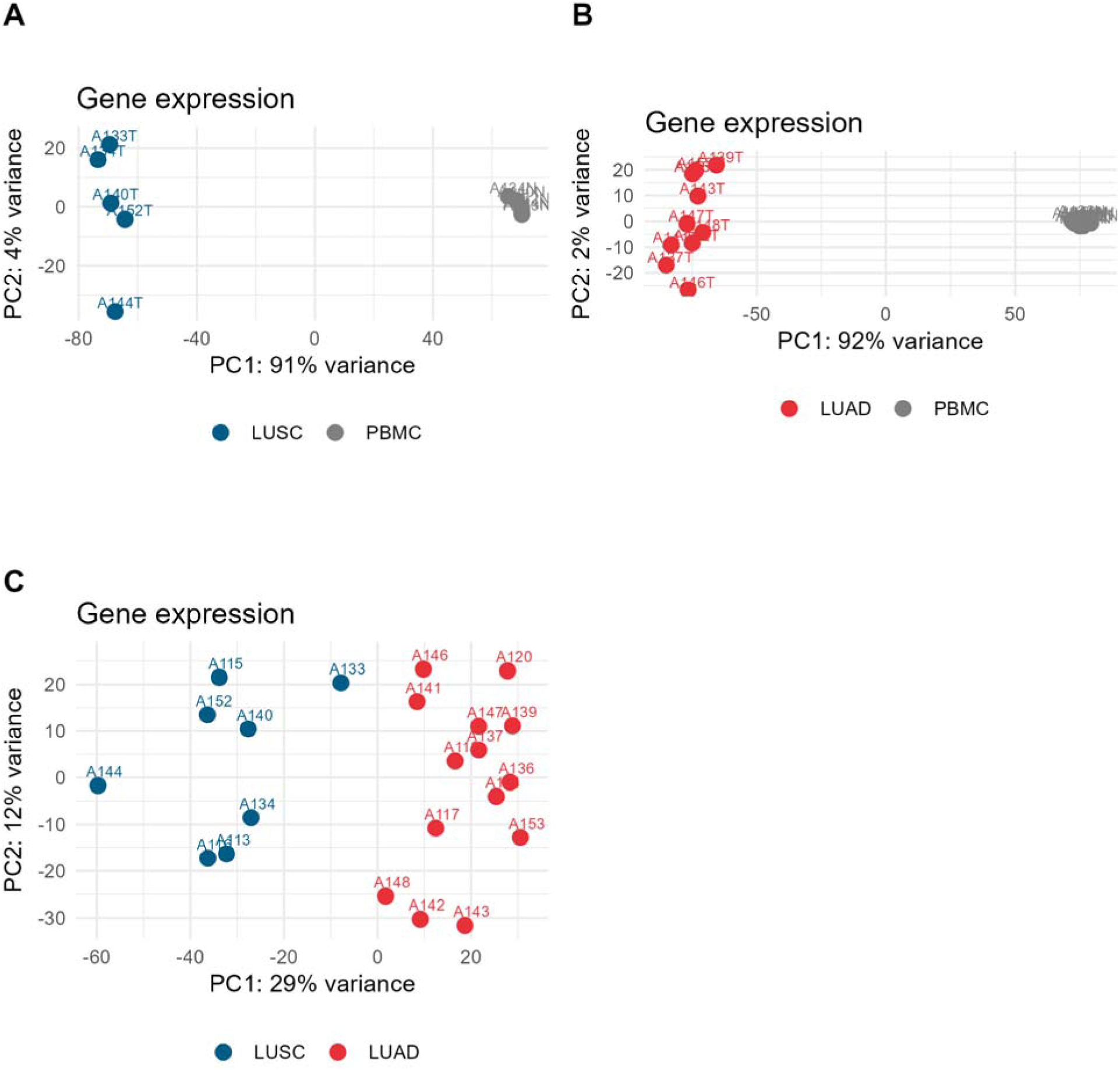
PCA of normalised gene count matrix. (A) LUSC (blue) & PBMC (grey). (B) LUAD (red) & PBMC (grey). (C) LUSC & LUAD. Samples are numbered with donor identifier.

Comparison between LUSC and LUAD found that the two cancer subtypes divide along PC1, but there are clusters within each sub-type and some individuals, notably LUSC individuals A144 and A133 (Figure 1 C). There were no obvious batch effects.The first three PCs account for 50% of the variance, and the first 10 PCs account for 88% of the variance (Figure S1).

Samples were grouped according to LUSC or LUAD subtype or PBMC and differential gene expression (DEG) was calculated from both genomic alignment and transcript classification count matrices using edgeR [16]. Each final DEG table was filtered for genes common to both analyses (Table 3). DEG for shared genes from genomic alignments with HISAT2 are shown here and the transcript classification results from Salmon are provided in the Supplementary Information.

**Table 3:** Comparison of DEGs.

| Comparison | Total DEGs | *LUSC DEGs | *LUAD DEGs |
| --- | --- | --- | --- |
| LUSC & PBMC | 17,719 | 3,989 | - |
| LUAD & PBMC | 17,586 | - | 3,822 |
| LUSC & LUAD | 19,859 | 265 | 51 |

\*DEG above thresholds of $\log_2$ fold-change of 1.5 and an FDR of 1%.

Table 3 shows the numbers of DEGs for each NSCLC subtype comparison exceeding thresholds of log_2_ fold-change of 1.5 and below a false discovery rate (FDR) of 1%. These thresholds are necessarily arbitrary and chosen to balance being conservative whilst not over-excluding information. The data without thresholds are provided in the Supplementary Information Tables S7-9.

When comparing NSCLC to PBMC we observe nearly 4,000 tumour DEGS for both subtypes (Table 3, Figure 2 A-B) . Whereas when comparing LUSC to LUAD only 316 of 19,859 DEGs exceeded thresholds of log_2_ fold-change of 1.5 and below a FDR of 1%. Of these 316 genes 265 were enriched in LUSC and 51 in LUAD (Table 3, Figure 2 C).

**Figure 2:**
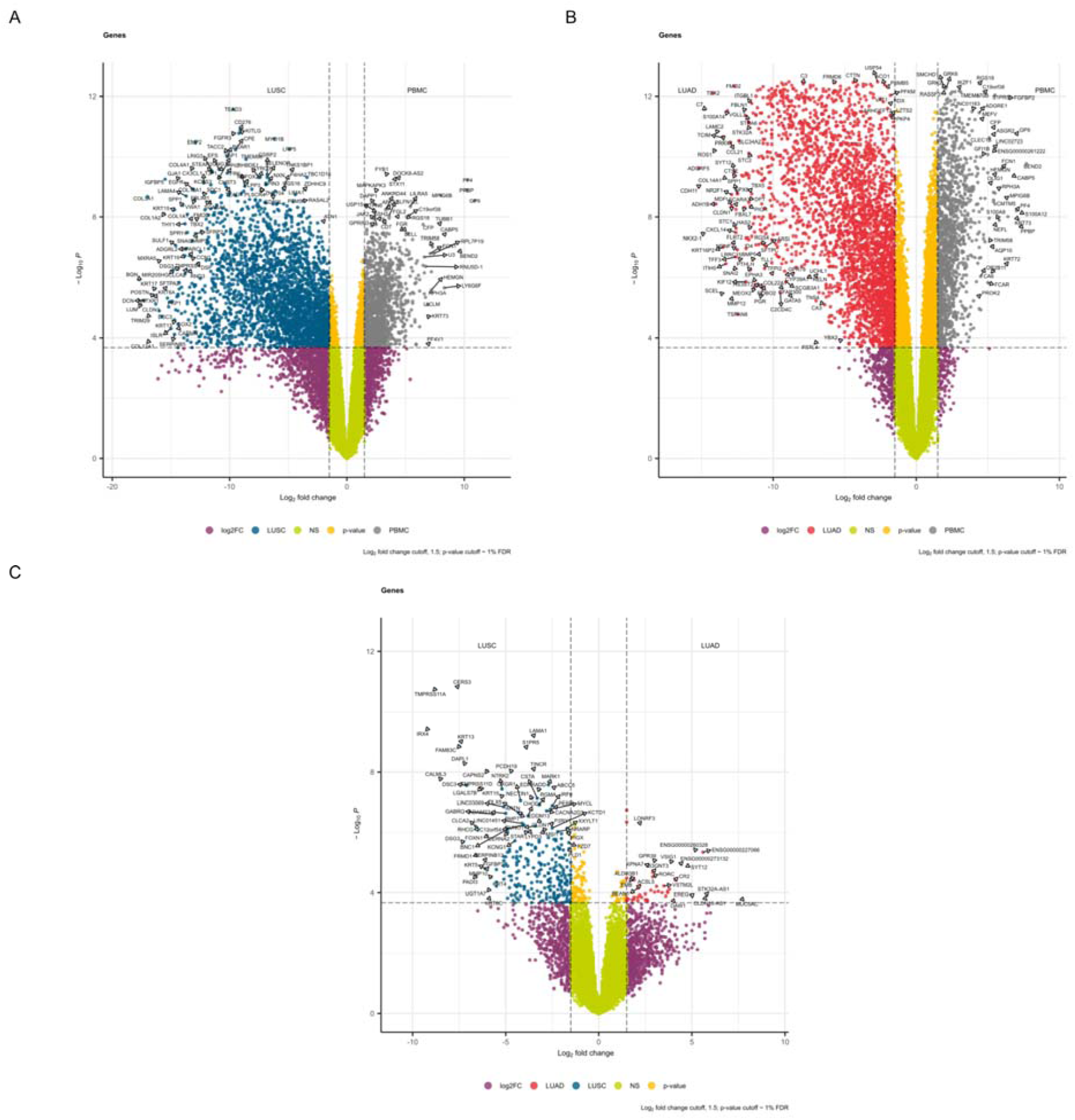
Volcano plots of DEGs. Thresholds are represented by dotted lines at FDR of 1% and log_2_ fold change of 1.5. (A) Comparison of LUSC & PBMC (n=17,719). (B) Comparison of LUAD & PBMC (n=17,586). (C) Comparison of LUSC & LUAD (n=19,859).

To further examine sample to sample variation of DEGs for each comparison, using just the significance threshold of 1% FDR, we plotted the log_2_ fold-change rescaled as z-scores i.e. each unit from zero represents one standard deviation from the row average value for each gene (Figure 3). These heatmaps reinforce the observations from Figure 1 of the high similarity between PBMC samples and dissimilarities between patient tumours, either when compared with PBMC (Figure 3 A-B) or between NSCLC subtypes (Figure 3 C). However, the overall dissimilarity between NSCLC and PBMC and between LUSC and LUAD in DEGs remains.

**Figure 3:**
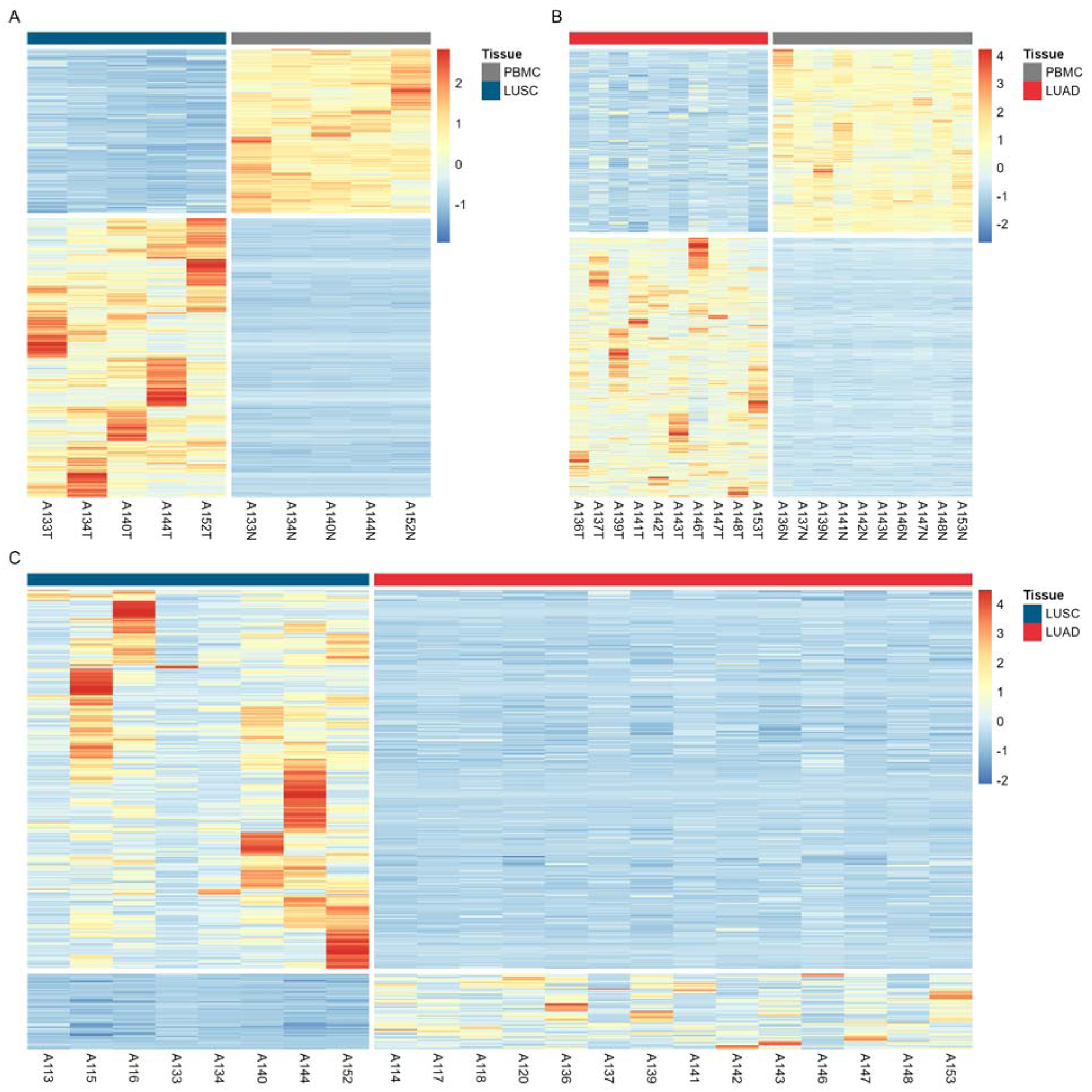
Heatmaps of DEGs below a FDR of 1%. Colour bar shows log_2_ fold change rescaled as z-scores i.e. each unit from zero represents one standard deviation from the row average value for each gene. Tumour samples are indicated with suffix T and PBMC samples with suffix N (A) Comparison of LUSC & PBMC (n=8,089). (B) Comparison of LUAD & PBMC (n=10,058). (C) Comparison of LUSC & LUAD (n=428).

### Comparison of proteomes

Proteins were quantified using label free quantification (LFQ) [17,18] yielding protein identifications from the normalised top 3 peptide intensities (Tables S10-12). As with the transcriptome data, we first performed PCA using the normalised top 3 peptide intensities of the 500 most variable proteins to examine separation between and within the conditions (Figure 4, Figure S2) [15]. There was clear separation along PC1 between NSCLC subtypes and NAT (Figure 4 A-B). However samples were dispersed along PC2 for both LUAD and NAT (Figure 4 A), with two NAT clusters for the LUSC comparison (Figure 4 A). This indicates homogeneity within NAT and heterogeneity between the tumours. The cancer subtype separation is much less clear for LUSC and LUAD comparison, PCs 1 and 2 account for around one third of the variance, and without the labels the division between them would not be obvious (Figure 4 C). LUAD patient A139 is an outlier, while LUSC patients A116, A140, A144 and A152 form a cluster.

**Figure 4:**
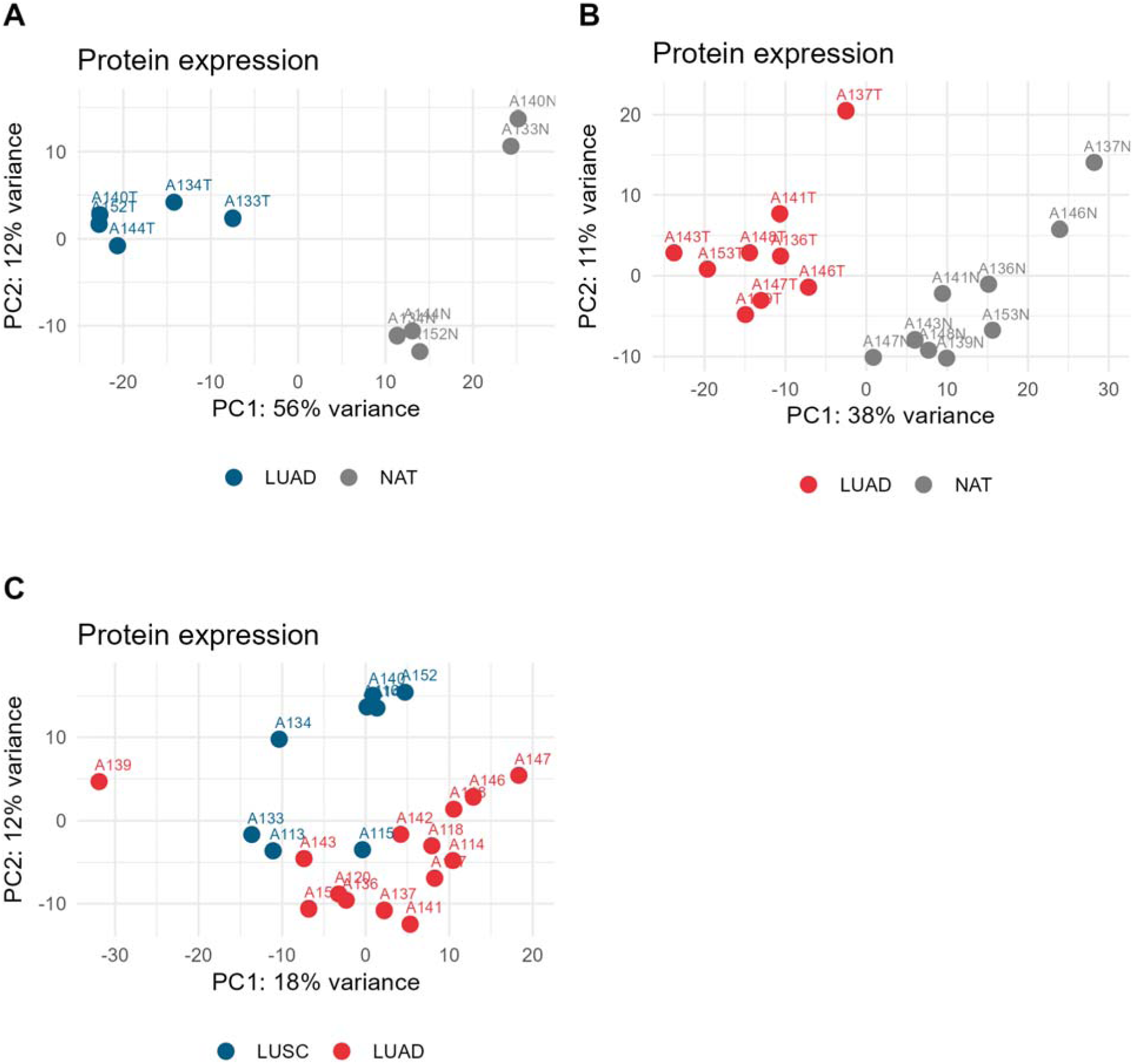
PCA of normalised top 3 peptide intensities of 500 most variable proteins. (A) LUSC (blue) & NAT (grey). (B) LUAD (red) & NAT (grey). (C) LUSC & LUAD. Samples are numbered with donor identifier.

Samples were grouped according to LUSC or LUAD subtype or NAT and differential protein expression was calculated with DEqMS [19]. Mass spectrometry proteomics quantifies far fewer proteins than transcriptomes do genes due to methodological differences. For any protein (or gene) to be analysed for differential expression, it must be present in all samples under consideration. The total number of DEPs quantified for NSCLC and NAT comparisons is approximately one third of the DEPs quantified comparing LUSC and LUAD (Table 4). This indicates that the NAT proteome is very different to that of tumour tissue. Conversely, the proteomes of the two NSCLC subtypes are similar to each other, as indicated in the PCA analysis (Figure 4 C).

**Table 4:** Comparison of DEPs.

| Comparison | Total DEPs | *LUSC DEPs | *LUAD DEPs |
| --- | --- | --- | --- |
| LUSC & NAT | 1,330 | 326 | - |
| LUAD & NAT | 1,478 | - | 185 |
| LUSC & LUAD | 3,872 | 117 | 15 |

\*DEP above thresholds of log<sub>2</sub> fold-change of 1 and a p-value of 1%.

For DEP we used more relaxed thresholds to filter the results than for the DEGs of a log_2_ fold-change of 1 and significance below a p-value of 0.01. As before, the data without thresholds is provided in the Supplementary Information Tables S13-16.

When comparing NSCLC to NAT we observed less than 1,500 tumour DEPs for both subtypes (Table 4, Figure 5 A-B). Comparing LUSC to NAT yielded 326 proteins enriched in LUSC (Figure 5 A), and comparing LUAD to NAT yielded 185 proteins enriched in LUAD (Figure 5 B). Whereas when comparing LUSC to LUAD only 132 of 3,872 DEPs exceeded the thresholds of log_2_ fold-change of 1 and below a p-value of 1%. Of these 132 proteins 117 were enriched in LUSC and 15 in LUAD (Table 4, Figure 5 C). Note that in Figure 5 gene names have been used for labels to make comparison with DEGs easier.

**Figure 5:**
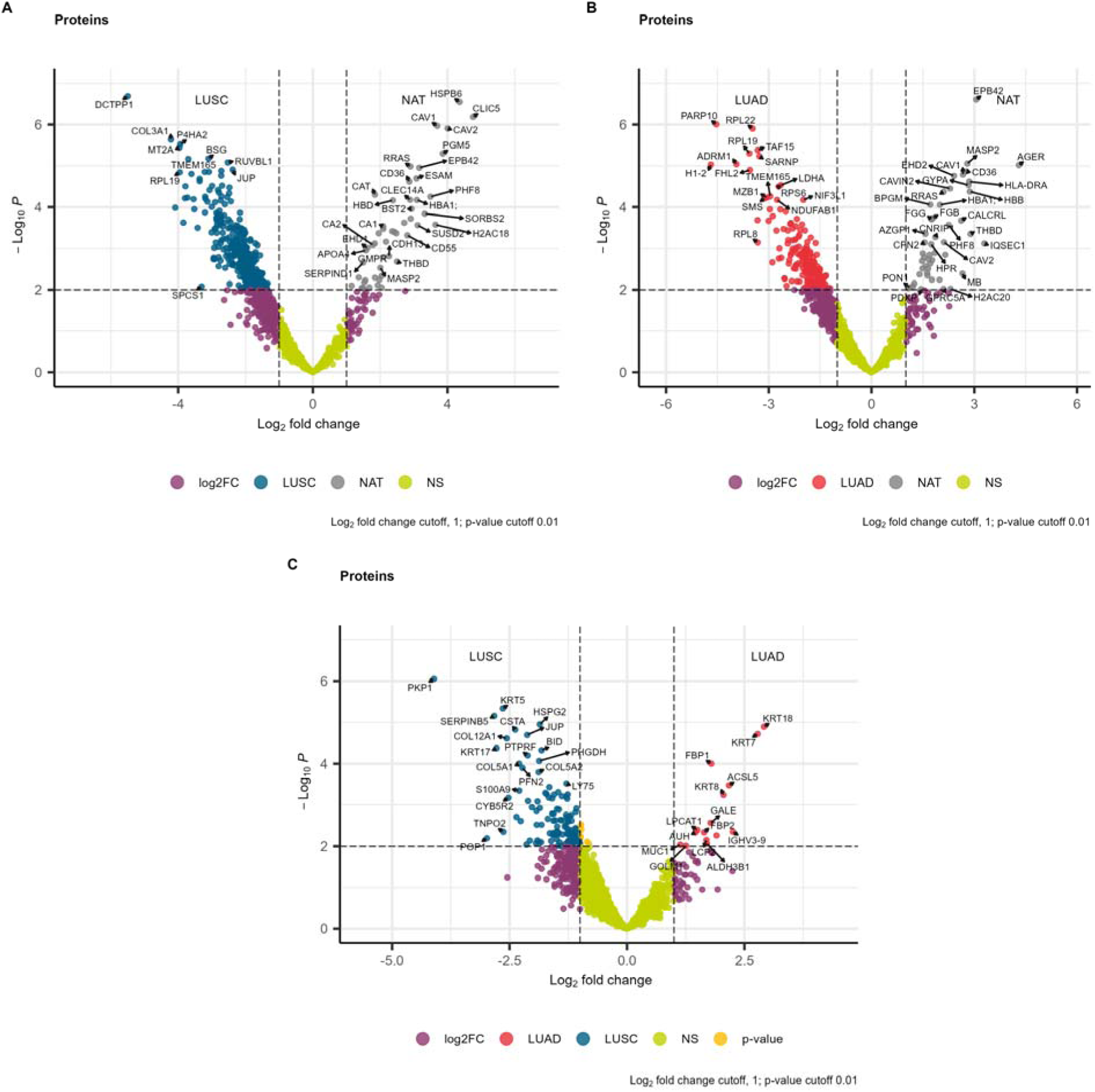
Volcano plots of DEPs. Proteins are labelled with gene names to ease comparison with DEG analysis. Thresholds are represented by dotted lines at p-value of 1% and log_2_ fold change of 1. (A) Comparison of LUSC & NAT (n=1,330). (B) Comparison of LUAD & NAT (n=1,478). (C) Comparison of LUSC & LUAD (n=3,872).

As with the transcriptomics, to further examine sample to sample variation of DEPs for each comparison, using just the significance threshold of below p-value of 1%, plotting the log_2_ fold-change rescaled as z-scores (Figure 6). Heterogeneity between tumour samples was again apparent for the comparisons with NAT, particularly for LUSC patients A133 and A140 (Figure 6 A), and LUAD A137 and A146 (Figure 6 B) corresponding with their separation from the main clusters on the PCA plots (Figure 4 A-B). Likewise we found the heterogeneity between tumour samples for the LUSC and LUAD comparison corresponds with spread of samples observed in the PCA plot (Figure 6 C). Stratification of the DEPs was not as clear as for the DEGs, but there were still a number of DEPs across all comparisons.

**Figure 6:**
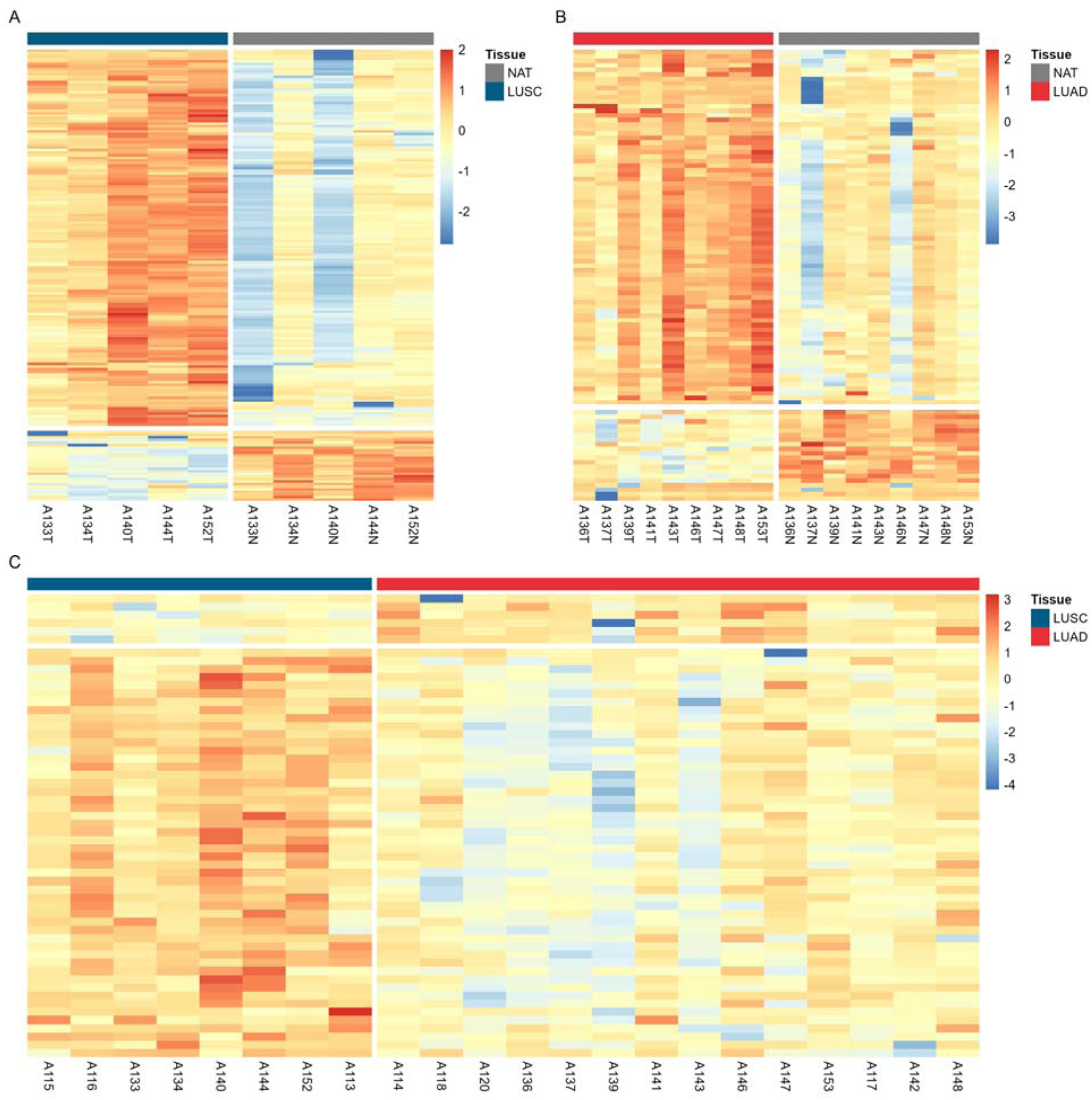
Heatmaps of DEPs below a p-value 1%. Colour bar shows log_2_ fold change rescaled as z-scores i.e. each unit from zero represents one standard deviation from the row average value for each protein. Tumour samples are indicated with suffix T and normal adjacent tissue with suffix N (A) Comparison of LUSC & NAT (n=379). (B) Comparison of LUAD & NAT (n=234). (C) Comparison of LUSC & LUAD (n=139).

## Functional analysis

Next we sought functional interpretations of the genes and proteins yielded from the differential expression analysis. We used g:Profiler to perform enrichment analysis to identify functional process and pathways [20]. To select the lists we chose thresholds that selected similar proportions of DEGs and DEPs from each comparison as inputs to g:Profiler. For the NSCLC and PBMC or NAT comparisons, DEGs and DEPs were filtered at and log_2_ fold-change of 1.5 and below 1% FDR. For between LUSC and LUAD comparisons DEGs were filtered at and log_2_ fold-change of 1.5 and below 5% FDR, whilst DEPs were filtered at and log_2_ fold-change of 1 and below a p-value of 5%. As previously, it is worth noting that the unfiltered DEGs and DEPs lists are provided in the Supplementary Materials Tables S7-9 and S13-16 for analysis with alternative thresholds.

g:Profiler performs enrichment analysis using eleven pathway sources and the full results are provided in the Supplementary Materials (Tables S17-24). Figure 7 and Figure 8 show up to the top 20 Gene Ontology Biological Process (GO:BP) pathways enriched for each comparison [21,22]. Figure 9 and Figure 10 show up to the top 20 Reactome pathways enriched for each comparison [23].

**Figure 7:**
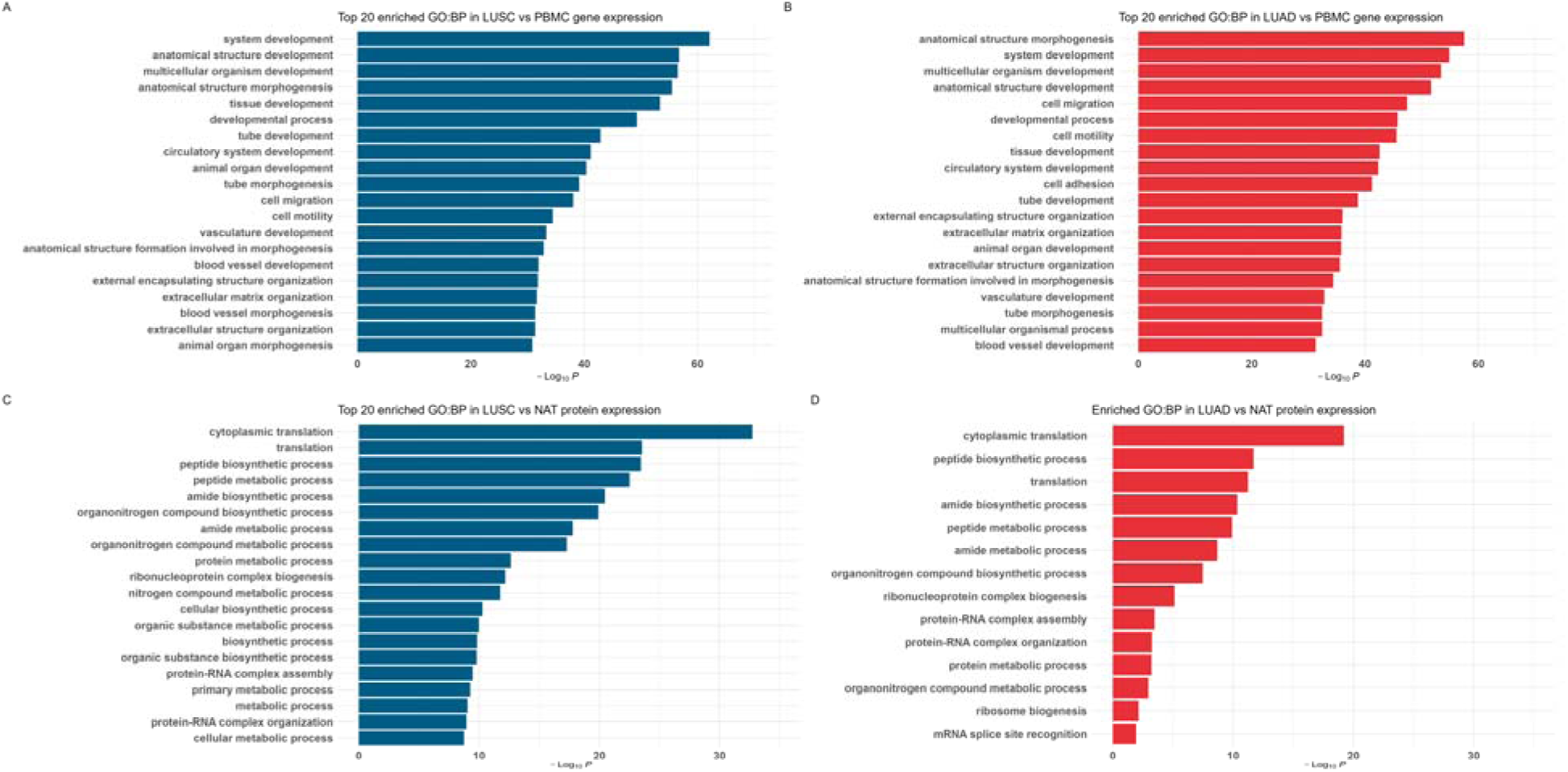
Bar plots of enrichment of GO Biological Processes of NSCLC subtypes vs PBMC or NAT. Up to the top 20 pathways are shown with statistical significance level indicated by the -log_10_ p-value on the x-axis. (A) Enriched in LUSC vs PBMC DEG (B) Enriched in LUAD vs PBMC DEG (C) Enriched in LUSC vs NAT DEP (D) Enriched in LUAD vs NAT DEP

**Figure 8:**
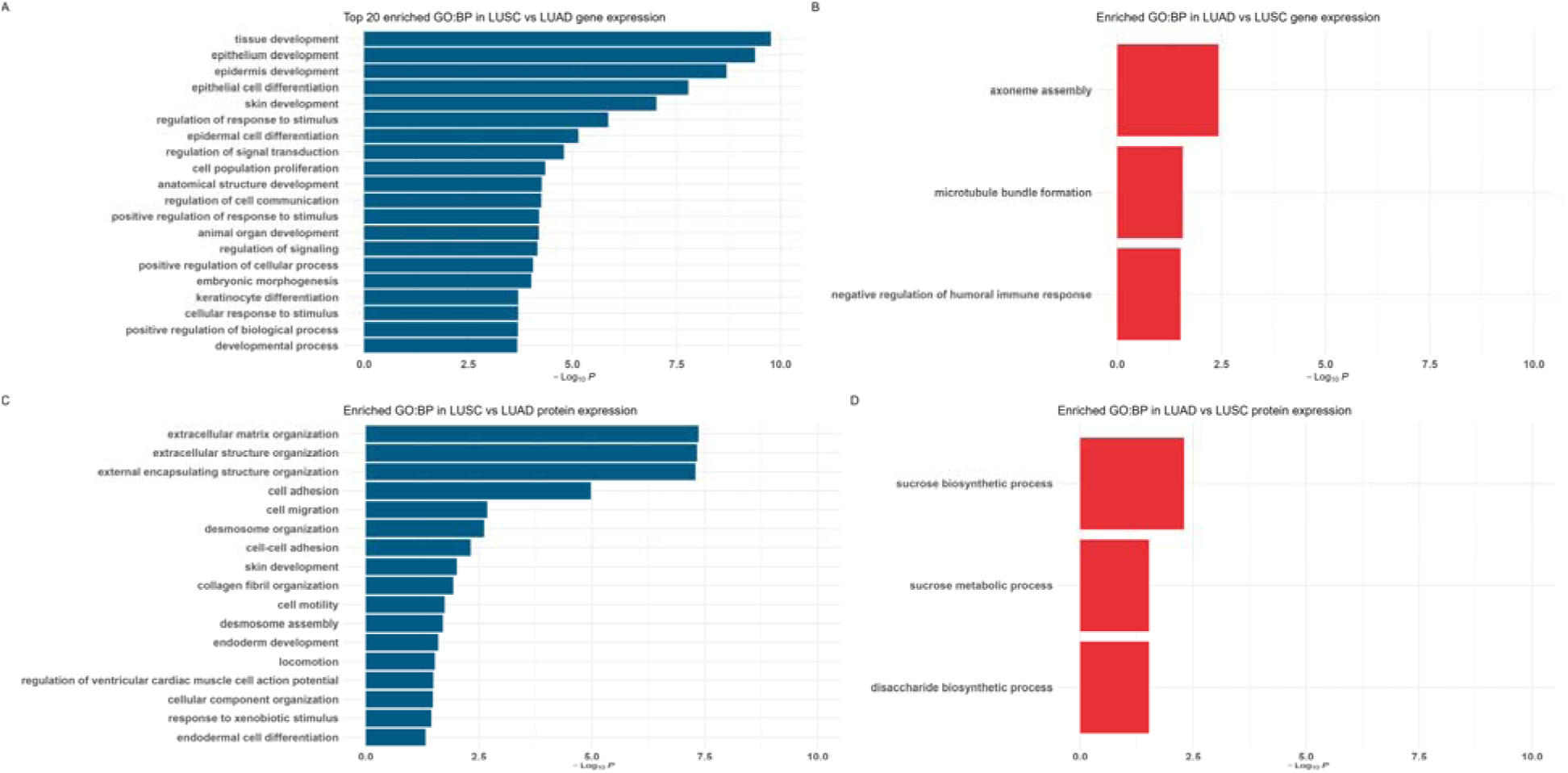
Bar plots of enrichment of GO Biological Processes comparing NSCLC subtypes. Up to the top 20 pathways are shown with statistical significance level indicated by the -log_10_ p-value on the x-axis. (A) Enriched in LUSC vs LUAD DEG (B) Enriched in LUAD vs LUSC DEG (C) Enriched in LUSC vs LUAD DEP (D) Enriched in LUAD vs LUSC DEP

**Figure 9:**
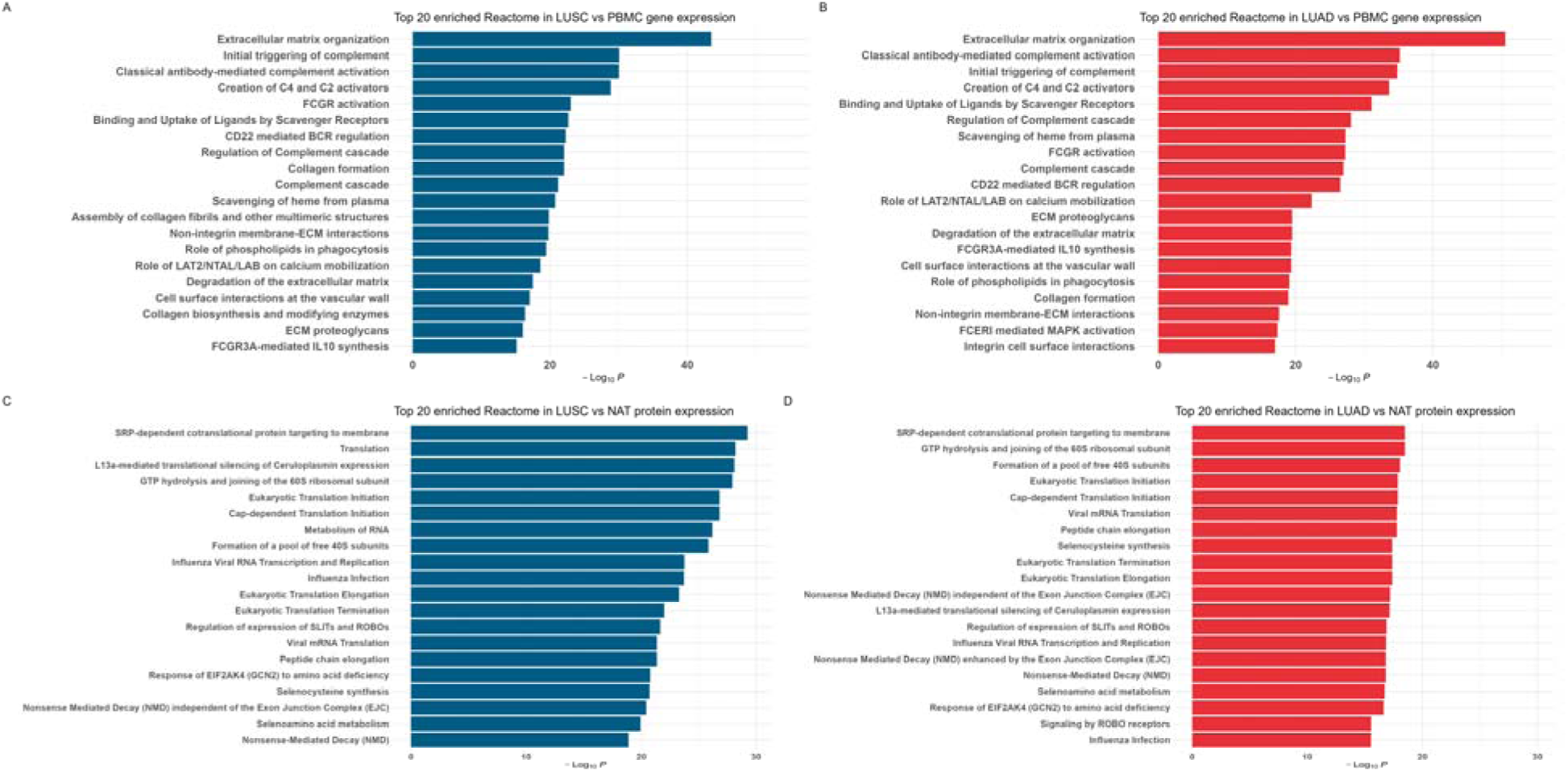
Bar plots of enrichment of Reactome pathways of NSCLC subtypes vs PBMC or NAT. Up to the top 20 pathways are shown with statistical significance level indicated by the -log_10_ p-value on the x-axis. plotted on the x-axis. (A) Enriched in LUSC vs PBMC DEG (B) Enriched in LUAD vs PBMC DEG (C) Enriched in LUSC vs NAT DEP (D) Enriched in LUAD vs NAT DEP

**Figure 10:**
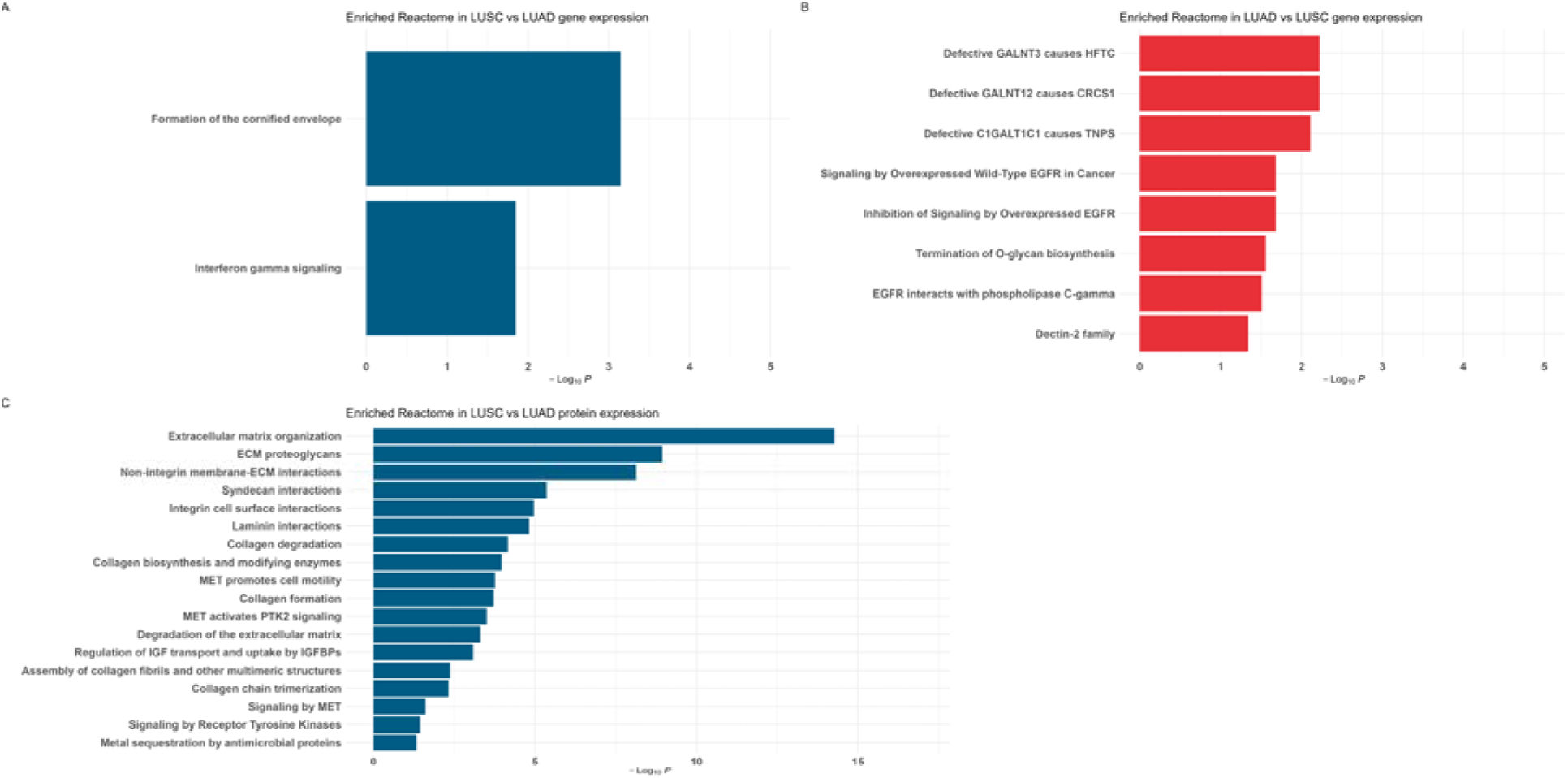
Bar plots of enrichment of Reactome pathways comparing NSCLC subtypes. Up to the top 20 pathways are shown with statistical significance level indicated by the - log_10_ p-value on the x-axis. plotted on the x-axis. (A) Enriched in LUSC vs LUAD DEG (B) Enriched in LUAD vs LUSC DEG (C) Enriched in LUSC vs LUAD DEP

### Enrichment of biological processes

GO:BP enrichment for comparisons of the NSCLC subtypes with PBMC indicates similarly enriched pathways for processes relating to developmental and structural changes for both subtypes (Figure 7 A-B). Comparisons of the NSCLC subtypes with NAT for GO:BP enrichment also identified similar pathways for both cancer subtypes, but this time relating to protein translation and RNA related processing and splicing (Figure 7 C-D).

Gene expression comparison between the NSCLC subtypes indicates enriched GO:BP for developmental processes for LUSC as seen for the PBMC comparison (Figure 8 A, Figure 7 A) . Whereas for LUAD, we identified three pathways relating to cellular structure, cell motility and immune evasion (Figure 8 B).

Protein expression comparison between the NSCLC subtypes indicates enriched GO:BP for extracellular matrix remodelling and cell migration and motility for LUSC, whereas metabolic processes including glucose metabolism are enriched in LUAD (Figure 8 C-D).

### Enrichment of Reactome pathways

Enrichment of Reactome pathways Gene expression for NSCLC subtypes compared to PBMC identified an identical set of pathways relating to extracellular matrix remodelling and immune system activation (Figure 9 A-B). Comparisons of the NSCLC subtypes with NAT correspond with the GO:BP enrichment for pathways we identified relating to protein synthesis, RNA processing and mRNA splicing (Figure 9 C-D).

Enrichment of Reactome pathways for gene expression of LUSC compared to LUAD identified two pathways, as for GO:BP, relating to extracellular matrix remodelling and immune system activation (Figure 10 A). Whereas for LUAD compared to LUSC identified Reactome pathway enrichment, as for GO:BP, relating to metabolic processes including altered glycosylation and immune system pathways (Figure 10 B).

No Reactome pathways were found enriched for protein expression of LUAD compared to LUSC, but for LUSC compared to LUAD extracellular matrix remodelling pathways were enriched (Figure 10 C).

## Discussion

Previously we used proteogenomics analysis to identify patient specific neoantigens arising NSCLC mutations as therapeutic targets [7]. Mutations in LUAD driver genes such as in epidermal growth factor receptor (EGFR) and ALK tyrosine kinase receptor (ALK) are also prognostic for therapeutic responses [24,25], whereas for LUSC driver mutations remain unknown. Whilst mutations have specific effects on genes, such as drug resistance in EGFR, collectively they also create chromosome instability and affect cell differentiation, cell cycle and epigenetic regulation, cell signalling and metabolism [26,27]. Here we identified evidence of collective mutational effects by examining differential transcriptomic and proteomic expression in two NSCLC subtypes, compared against each other, and against PBMC or NAT.

The main limitations in our design were that we were unable to compare NSCLC transcriptomes to NAT transcriptomes, and that we examined a relatively small cohort. Use of PBMCs will have confounded our observations to some extent, particularly with respect to expression of keratins and other genes that phenotypically differentiate lung tissues from blood. Hence this limits interpretation of the NSCLC and PBMC DEG comparison, but not the NSCLC subtype DEG comparison.

Two unsurprising observations were that tumour tissue transcriptomes or proteomes do not resemble those of either PBMC or NAT respectively, nor do NSCLC subtypes resemble each other. We were able to separate the groups each comparison by means of PCA of their gene counts and protein peptide intensities. We then examined differential expression of genes and proteins to characterise each comparison.

For both LUAD and LUSC we observed many similar DEGs between tumour and PBMC relating to: extracellular matrix remodelling and cell structure with DEG of several collagen and keratin genes such as COL1A1 and KRT19; cell adhesion, growth factors and signalling such as pulmonary surfactant-associated protein A2 (SFTPA2), EGFR, vascular endothelial growth factor receptor 2 (KDR) and proto-oncogene tyrosine-protein kinase ROS (ROS1); and cell differentiation, such as transcription factor SOX2, neurogenic locus notch homolog protein 3 (NOTCH3) and Tumour protein 63 (TP63) (Table S7-S8). These findings are consistent with previous observations for NSCLC [27,28] and are reflected in our functional analysis. Pulmonary surfactant SFTPA2 and its associated mutations were recently identified for use as a serum biomarker of NSCLC [4]. This is suggestive of common processes in tumourigenesis and potentially also clonality in tumour tissue versus heterogeneous normal tissue. As well as DEGs, we identified some of the same SFTPA2 mutations in several of our donors [7].

Metabolic processing genes Indoleamine 2,3-dioxygenase 1 (IDO1) and fatty acid synthase (FASN) were also both DEGs in both NSCLC subtypes relative to PBMCs and for which targeting drugs are either approved or in clinical trials [29].

We found two notable differences in gene expression in relation to immune inhibition between NSCLC subtypes as previously observed [30]: fibrinogen-like Protein 1 (FGL1) was a DEG in LUAD relative to LUSC. FGL1 has been identified as a T-cell suppressor as a ligand of LAG-3 [31]. Whereas autoimmune checkpoint gene V-set domain-containing T-cell activation (VTCN1) [32] was a DEG in LUSC relative to LUAD. These observations for FGL1 in LUAD and VTCN1 in LUSC support their potential as subtype specific targets for checkpoint inhibitor drugs (Tables S9 and S15).

As expected, between NSCLC subtypes Napsin-A (NAPSA) was both a DEG and DEP in LUAD relative to LUSC, and DEG of Thyroid transcription factor 1 (NKX2-1), supporting their known utility as immunohistological LUAD classifiers (Tables S9 and S15) [33]. A machine learning NSCLC classification model of DEPs identified, in addition to NAPSA, identified DEP of Anterior gradient protein 3 (AGR3) as a feature of LUAD, and DEP of KRT5 and SERPINB5 as features of LUSC that were also present in our observations [30] (Table S15).

Ribosomal proteins, such as small ribosomal subunit proteins eS19 and uS10 (RPS19, RPS20), were differentially expressed between both NSCLC subtypes and NAT. These and other ribosomal proteins are implicated in regulation of TP53 [34] and are indicative of the changes between tumour and normal tissue seen in the functional analysis identification of pathways relating to protein translation and RNA-related processes (Figures 7-8, Table S13-S14).

Proteins in LUSC differentially expressed in comparison to LUAD included Poly ADP-ribose polymerase 1 (PARP1) a target for the DNA repair inhibitor Olaparib. Results for PARP1 inhibition in a recent NSCLC trial for patients with homologous repair deficiency were inconclusive [35], but another trial is ongoing (NCT03976362). Epigenome histone deacetylases (HDAC1, HDAC2) were also DEP for LUSC and is under trial as a target for entinostat as an inhibitor/chemosensitizer (NCT05053971) [36,37]. DEP RAC-alpha serine/threonine-protein kinase (AKT1) has been identified playing a role transdifferentiation of LUAD to LUSC [38].

DEPs Transferrin receptor protein 1 (TFRC) and Phosphoserine aminotransferase (PSAT1) have been previously identified as characteristic of a LUSC subtype relating to changes in metabolic signalling and oxidative stress [39].

For LUAD, DEPs compared to LUSC included: Mucin-1 (MUC1) a target for salinomycin [40], Serine/threonine-protein kinase mTOR (MTOR) which has been identified as a chemosensitizing target [41], whilst enzymes Transglutaminase 2 (TGM2) and Sterol O-acyltransferase 1 (SOAT1) have been identified as targets for inhibition [42,43] (Table S15). An intriguing LUAD DEP is B-cell lymphoma/leukemia 10 (BCL10) which has a role in inflammation as part of the multi-protein complex in the NF-LB pathway [44,45] and therefore may relate to activation of EGFR [46]. In B-cell lymphomas, trials for inhibitors targeting another protein in the same complex, Mucosa-associated lymphoid tissue protein 1 (MALT1), are ongoing, as well as efforts to understand their efficacy in solid cancers [47].

Overall our observations demonstrate that comparably high quality data can be produced from a modestly sized cohort. They show good agreement with published NSCLC proteogenomics studies and provide confirmatory evidence for current pharmacological intervention studies.

## Materials and Methods

### Ethics statement

Ethical approval was obtained from the local research ethics committee (LREC reference 14-SC-0186 150975) and written informed consent was provided by the patients.

### Tissue preparation

Tumours were excised from resected lung tissue post-operatively by pathologists and processed either for histological evaluation of tumour type and stage, or snap frozen at −80°C. Whole blood samples were obtained, and PBMCs were isolated by density gradient centrifugation over Lymphoprep prior to storage at −80°C.

### RNA extraction

RNA was extracted from tumour tissue that had been obtained fresh and immediately snap frozen in liquid nitrogen. Ten to twenty 10 µm cryosections were used for nucleic acid extraction using the automated Maxwell® RSC instrument (Promega) and Maxwell RSC simplyRNA tissue kit according to the manufacturer’s instructions. RNA was quantified using Qubit fluorometric quantitation assay (ThermoFisher Scientific) according to the manufacturer’s instructions. RNA quality was assessed using the Agilent 2100 Bioanalyzer generating an RNA integrity number (RIN; Agilent Technologies UK Ltd.).

### RNA sequencing

Samples were prepared as TruSeq stranded mRNA libraries (Illumina, San Diego, USA) and 100bp paired end sequencing was performed using the Illumina NovaSeq 6000 system by Edinburgh Genomics (Edinburgh, UK). Raw reads were pre-processed using fastp (version 0.20.0) [48].

Filtered reads were aligned twice: Firsty to the 1000 genomes project version of the human genome reference sequence (GRCh38/hg38) using HISAT2 (version 2.2.1) [12], merged and then transcripts assembled and gene expression estimated with featureCounts (version 2.0.6) [13] using reference guided assembly. Secondly reads were aligned and quantified using transcript classification with Salmon (version 1.10.3) [14].

### Differential gene expression

Differentially expressed genes (DEGs) were estimated using transcript counts from both HISAT2 and salmon using edgeR using default settings [16]. The intersection of DEGs common to both the HISAT2 and salmon analysis were used to filter the HISAT2 results that were used for the remaining analysis.

Principal component analysis of the normalised HISAT2 count matrices was performed using DESEq2 [15] and PCATools [49].

Results were visualised using EnhancedVolcano [50], pheatmap [51] and ggplot2 [52].

### Protein extraction and digestion

Snap frozen tissue samples were briefly thawed and weighed prior to 30s of mechanical homogenization (Fisherbrand homogenizer 150 using plastic generator probes, Fisher Scientific, UK), in 8 mL lysis buffer (0.02 M Tris, 0.5% (w/v) IGEPAL, 0.25% (w/v) sodium deoxycholate, 0.15 mM NaCl, 1 mM EDTA, 0.2 mM iodoacetamide supplemented with EDTA-free protease inhibitor mix) and incubated at 4°C for 30 min. Homogenates were then centrifuged at 2,000 g for 10 min at 4°C to remove cell debris, and for a further 60 min at 13,000 g, 4°C to clarify. Supernatant was stored at -80°C prior to protein extraction for proteomic analysis.

Protein concentration of tissue lysates was determined by BCA assay, and volumes equivalent to 100 mg of protein were precipitated using methanol/chloroform as previously described [53]. Pellets were briefly air-dried prior to resuspension in 6 M urea/50 mM Tris-HCl (pH 8.0).

Proteins were reduced by the addition of 5 mM (final concentration) DTT and incubated at 37°C for 30 min, then alkylated by the addition of 15 mM (final concentration) iodoacetamide and incubated in the dark for 30 min. 4 µg Trypsin/LysC mix (Promega) were added and the sample incubated for 4 h at 37°C, then 6 volumes of 50 mM Tris-HCl pH 8.0 were added to dilute the urea to < 1 M, and the sample was incubated for a further 16 h at 37°C. Digestion was terminated by the addition of 4 µL of TFA, and the sample clarified at 13,000 x g for 10 min at RT. The supernatant was collected and applied to Oasis Prime microelution HLB 96-well plates (Waters, UK) which had been pre-equilibrated with acetonitrile. Peptides were eluted with 50 µL of 70% acetonitrile and dried by vacuum centrifugation prior to resuspension in 0.1% formic acid.

### Mass spectrometry proteomics

8 µg of peptides per sample were separated by an Ultimate 3000 RSLC nano system (Thermo Scientific) using a PepMap C18 EASY-Spray LC column, 2 µm particle size, 75 µm x 75 cm column (Thermo Scientific) in buffer A (H_2_O/0.1% Formic acid) and coupled on-line to an Orbitrap Fusion Tribrid Mass Spectrometer (Thermo Fisher Scientific,UK) with a nano-electrospray ion source.

Peptides were eluted with a linear gradient of 3-30% buffer B (acetonitrile/0.1% formic acid) at a flow rate of 300 µL/min over 200 min. Full scans were acquired in the Orbitrap analyser in the scan range 300-1,500 m/z using the top speed data dependent mode, performing an MS scan every 3 second cycle, followed by higher energy collision-induced dissociation (HCD) MS/MS scans. MS spectra were acquired at a resolution of 120,000, RF lens 60% and an automatic gain control (AGC) ion target value of 4.0e5 for a maximum of 100 ms. MS/MS scans were performed in the ion trap, higher energy collisional dissociation (HCD) fragmentation was induced at an energy setting of 32% and an AGC ion target value of 5.0e3.

### Proteomic data analysis

Raw spectrum files were analysed using Peaks Studio 10.0 build 20190129 [17,54] and the data processed to generate reduced charge state and deisotoped precursor and associated product ion peak lists which were searched against the UniProt database (20,350 entries, 2020-04-07) plus the corresponding mutanome for each sample (∼1,000-5,000 sequences) and contaminants list in unspecific digest mode. Parent mass error tolerance was set a 10ppm and fragment mass error tolerance at 0.6 Da. Variable modifications were set for N-term acetylation (42.01 Da), methionine oxidation (15.99 Da), carboxyamidomethylation (57.02 Da) of cysteine. A maximum of three variable modifications per peptide was set. The false discovery rate (FDR) was estimated with decoy-fusion database searches [17] and were filtered to 1% FDR.

### Differential protein expression

Label free quantification using the Peaks Q module of Peaks Studio [17,18] yielding matrices of protein identifications as quantified by their normalised top 3 peptide intensities. The resulting matrices were filtered to remove any proteins for which there were more than two missing values across the samples. Differential protein expression was then calculated with DEqMS using the default parameters [19].

Principal component analysis of the normalised top 3 peptide intensities was performed using DESEq2 [15] and PCATools [49].

Results were visualised using EnhancedVolcano [50], pheatmap [51] and ggplot2 [52].

### Functional analysis

Functional enrichment analysis was performed using g:Profiler [20] using default settings for homo sapiens modified to exclude GO electronic annotations. Gene ids were used as inputs for DEGs and protein ids for DEPs.

### Data availability

RNAseq data have been deposited at the European Genome-phenome Archive (EGA) under EGA Study ID: EGAS00001005499

The mass spectrometry proteomics data have been deposited to the ProteomeXchange Consortium via the PRIDE [55] partner repository with the dataset identifier PXD054390 and 10.6019/PXD054390.

PRIDE Reviewer account details: Username: Password: 364llR3qsAIS

Supplementary Figures S1 and S2 and Tables S1-S17 are available on Github: https://github.com/ab604/lung-global-supplement and Zenodo DOI: 10.5281/zenodo.13327662

## Acknowledgements

This study was supported by a Cancer Research UK Centres Network Accelerator Award Grant (A21998). Instrumentation in the Centre for Proteomic Research is supported by the Biotechnology and Biological Sciences Research Council, Grant/Award Number: BM/M012387/1.

## Notes

https://github.com/ab604/lung-global-supplement

https://zenodo.org/doi/10.5281/zenodo.13327662

### Competing Interest Statement

The authors have declared no competing interest.

